# Intra-individual variability in task performance after cognitive training is associated with long-term outcomes in children

**DOI:** 10.1101/2020.11.19.390427

**Authors:** A Cubillo, H Hermes, E Berger, K Winkel, D Schunk, E Fehr, T. A. Hare

## Abstract

The benefits and mechanistic effects of working memory training in children are the subject of much research and debate. The cumulative evidence indicates that training can alter brain structure and function in the short term and have lasting effects on behaviour. We show that five weeks of working memory training led to greater activity in prefrontal and striatal brain regions, better accuracy, and reduced intra-individual variability in response times. The reduction in intra-individual variability can be explained by changes to the evidence accumulation rates and thresholds in a sequential sampling decision model. Critically, intra-individual variability was more closely associated with academic skills and mental health 6-12 months after the end of training than task accuracy. These results indicate that intra-individual variability may be a useful way to quantify the immediate impact of cognitive training interventions and predict the future emergence of academic and socioemotional skills.

## Introduction

Cognitive training programs have received considerable attention over the years given their potential to improve cognitive abilities in healthy and clinical populations. However, the effectiveness and persistence of benefits from cognitive training programs are still being closely examined and vigorously debated^1-9^. Although cognitive training programs have been shown to improve performance on similar untrained tasks (near transfer), the evidence for transfer to cognitive skills in other domains (far transfer) remains more sparse and controversial^1-5,8,9^. We still lack sufficient understanding of the types of cognitive skills and abilities that are most beneficial to train, types of training methods and dosages that work best for particular skills, and the types of individuals that can reap sufficient benefits to justify the time and monetary costs of cognitive training interventions.

A recent study of over 500 first-grade children has generated important new findings indicating that the effects of training can emerge and increase over time^10^. This study found that the far-transfer benefits from adaptive working memory training to academic skills (e.g. Reading and Geometry) were only evident 6-12 months after the end of training. Moreover, this work showed that five weeks of adaptive working memory training during the first-grade year led to an increased probability of entering the highest academic track of the German secondary school system 3-4 years later. Given these results, longitudinal study designs that include follow-up measures over multiple years will be important for determining the potential effectiveness of different types and/or doses of cognitive training, especially for children.

It is important to understand the cognitive and neurobiological changes that take place during or just after training. Presumably, these proximal effects allow for the eventual emergence of wider-ranging benefits in the future. Cognitive training that resulted in near- or far-transfer effects has been reported to alter brain structure and function^11-16^. Training induced changes are often observed in prefrontal, parietal, and striatal regions that support executive functions such as working memory and attention control^17-21^. Here, we will define attention control as the cognitive processes required to coordinate and allocate attention to the relevant stimuli in the environment^22^. Cross sectional and longitudinal studies suggest that the maturation of these brain regions supports improvements in task performance over the course of development from childhood into adulthood^23-28^, and deterioration in these regions and their connectivity is associated with cognitive decline during aging^29-31^. Brain imaging studies have also indicated that successful transfer from trained to untrained skills requires that both cognitive processes engage at least partially overlapping structural and functional brain systems^32,33^. Thus, ideally, cognitive training programs should facilitate neural developments, potentially including maturation processes in children, that allow for more effective and efficient engagement of both specialized and domain general brain functions. This is because, by definition, domain general brain functions will overlap with the processes that support and facilitate a broader range of untrained skills.

Sensitive and reliable measures of changes in mental functions are necessary to detect the immediate effects of training interventions and forecast their long-term benefits. Here, we test the hypothesis that intra-individual variability in task performance – quantified via response times – can be used to assess training efficacy in the short term and is correlated with future far-transfer effects. Intra-individual variability measures derived from response times are well suited to this purpose because they have good test-retest reliability when computed across a sufficient number of trials, and are as, or even more, sensitive and robust indicators of cognitive function than the mean or variance in accuracy^34,35^.

Intra-individual variability in performance is associated with the cognitive abilities and brain functions targeted by cognitive training interventions in healthy individuals as well as individuals with various psychiatric and neurological conditions^36-38^. Developmentally, intra-individual variability shows an inverted-U shaped association with age, decreasing (i.e. improving) from childhood through late adolescence until young adulthood, and then increasing (i.e. worsening) again in old age^28,39,40^. Thus, intra-individual variability tracks the well-established pattern of brain and cognitive development across the lifespan. Therefore, intra-individual variability could plausibly detect changes in these systems caused by training interventions. Intra-individual variability is associated with the integrity of brain structure and function, most strongly in frontal brain regions, as well as with dopaminergic neuromodulation see ^34 for a review,38,40-44^. Furthermore, intra-individual variability measures are closely associated with the inhibitory and cognitive control abilities mediated by frontal and subcortical dopaminergic brain systems in healthy children and adults^28,42,45-47^. In summary, there is sufficient reason to hypothesize that intra-individual response time variability metrics can detect short-term training effects and may be useful in predicting the degree of long-term benefits.

We use a combination of cognitive tasks (N-Back and Flanker), functional magnetic resonance imaging (fMRI), and computational modelling of individual performance to examine the effects of five weeks of adaptive working memory training on brain and cognitive function in first-grade children. Working memory training improved response accuracy and decreased intra-individual variability on both tasks. Moreover, we show that the improvements in intra-individual variability detected during N-Back and Flanker task performance right after training are associated with the children’s academic skills and level of behavioural and psychological adjustment 6-12 months after training was completed.

## Methods

### Participants

Twenty-eight typically developing 7 to 9 year-old primary school children (mean age 7y9m, SD 5m, 14 females, working memory training = 16, comparison group = 12) participated in the study. These children were recruited out of an ongoing intervention study of over 500 children and 29 different classrooms. All participants had normal or corrected-to-normal vision and all but 2 were right-handed. No present or past psychiatric diagnosis were reported. Three of the participants attended schools where, unbeknown to the experimenters at the time of data collection, the randomization process was compromised. Specifically, the school authorities agreed to take part in the study only if their classrooms were included in the “instruction as usual” control group. The children and their families were still blind to the treatment condition that their classroom had been allocated to. We ran robustness checks for all analyses that excluded those 3 children and found similar results in all cases. The local ethics committee (Kantonale Ethikkommission Zürich) approved all procedures and methods used during this study. Participants received compensation for the participation in the study as described below.

### Recruitment and general procedure

Parents were informed of the upcoming study either during a regularly scheduled Parents’ Evening event taking place at school or in a specific meeting about the study. They were given a leaflet with the contact details for the team and those who would be willing to know more about the study or having interest in their children taking part in the study. Once they contacted the team, they were given an appointment with one of the research assistants at a local school, so that they could ask any questions and the children would have information about what the fMRI session would look like and what type of tasks they would have to perform while inside the scanner. To help them get familiar with how enclosed they would be and the scanner noise they were asked to lie flat in a play tunnel for approximately 6 minutes. During this time, they would see a movie of the different tasks and heard the scanner noise. If they agreed to take part, they would receive an appointment for the scanner session. On the day of the session, they were reminded again of the different tasks that they would perform while inside the scanner, and had the chance to practice them on a laptop until it was clear they understood and were able to perform those tasks correctly.

Next, a big teddy bear was used to show them the positioning procedures in the scanner, in order to reduce their potential anxiety and to have the chance to ask any questions they might have. Finally, once they had been positioned, they could ask for the accompanying parent to stay in the scanner room with them until they felt safe.

For participating in the fMRI portion of the study, the parents were compensated for the travel expenses and time. Children received family day passes for the Zurich Zoo and depending on their performance in one of the tasks, tokens that could be exchanged for prizes (see Intertemporal Choice Task description in section ***Post-training cognitive and decision tasks*** below).

### Cognitive training program description

The training procedures consisted of a five-week intervention and four assessment waves, one pre-intervention (baseline – w1), one immediately after the end of the five-week intervention (w2), and two follow-up waves at 6 and 12-13 months respectively (w3 and w4). The assessment sessions were run by a professional data collection service. These sessions were conducted by interviewers specifically trained and recruited for the study who were experienced with standard procedures in this population and age group. The assessment battery included tests of working memory and IQ (digit span, location span, object span, Raven’s test), educational outcomes (math numeracy and math geometry, reading abilities) and concentration tests (Go/NoGo and bp task).

The study was conducted using a between-class design, that is, classes were randomly assigned to the different treatment options. Randomization was stratified based on SES variables including the proportion of low household incomes, social benefit recipients, and non-Swiss residents.

The working training program implemented was Cogmed’s RoboMemo^1^. It is a computerized program, highly adaptive to individual performance, implemented via notebook computers including headphones for the spoken instructions and an external mouse. These specially dedicated notebooks were distributed to the participating schools. The intervention consisted of a daily working memory training session per day (duration ∼ 30 min), over a period of 5 weeks (25 sessions in total). While the standard intervention consists of 13 different tasks, 3 of them include either letters or syllables that are not suitable for first graders and were therefore excluded. Most of the remaining tasks focus heavily on visuo-spatial working memory, and only two of them on verbal working memory (digit span tasks). While 5 of the tasks are simple span tasks, the remaining 5 can be considered complex working memory tasks, which involve some processing of the information. Each training session included 6 adaptive modules (working memory tasks), including each 12 trials (75 trials in total). During the intervention, there was one specifically trained student coach in each class.

We compare the working memory training group to children that either received standard classroom instruction or a five-week self-regulation training. The self-regulation training took place during school lessons once a week. In these lessons, the teacher taught a version of the mental contrasting with implementation intentions (MCII) technique^48^ that was adapted to the relevant age group and the classroom context.

### Post-training cognitive and decision tasks

Subjects practiced each task right before the scanning session. There were 20, 44, and 8 practice trials for the Flanker, N-Back, and intertemporal choice tasks, respectively.

#### Working memory (N-Back) task

The 11-min block design working memory task consists of four conditions (Figure S1a). Participants are presented with series of pictures (1s duration, followed by 1s fixation cross). In the ‘0-Back’ condition, they have to respond whenever they see the picture of a sun on the screen. In the ‘1-Back’, ‘2-Back’, and ‘3-Back’ conditions, they have to respond whenever the picture on the screen is the same as 1, 2, or 3 before it, respectively. They perform the task in 2 runs (∼5 min each). Each run consists of 8 pseudo-randomised blocks, with 12 stimuli (3 targets and 9 non-targets). Each condition is presented twice per run. Before each block they saw a green fixation cross (12s) followed by a black fixation cross (1s) to indicate they should be alert as instructions were coming. They were then shown instructions (5s) on which condition was coming next. Performance data were recorded during scanning. The main performance variables are percentage of correct responses (correctly identified targets/total number of targets), percentage of commission errors (number incorrect responses/total number of non-target stimuli), mean reaction times to correctly identified targets, and variability in response times for correct responses.

#### Flanker task

The 11-min event-related task was designed based on^49^. Participants were presented with 240 trials shown in 2 separate runs (∼5 min duration each, Figure S1b). Each trial consisted in a central row of 5 yellow fishes over a blue background. They were instructed to “feed” the fish that was located in the centre of the screen. To do so, the child had to press the right/left button on the button box, depending on the direction of the central fish and ignoring the direction of the flankers, which on congruent/incongruent trials point towards the same/opposite direction, respectively. Each trial consisted of a 1s fixation cross, followed by the stimulus (1s), and finishing with a blank screen (random duration 0.1-1.5 s). The main performance variables are the % of correct responses, % of incorrect responses, reaction times, and variability of response times for each trial type (congruent and incongruent).

#### Intertemporal Choice Task (ICT)

We used an intertemporal choice task similar to one in ^50^. The task consisted of 48 trials, presented in 2 separate rounds, lasting a total of 12 minutes (Figure S1c). Each trial consisted of a fixation cross (duration random 3-6s) followed by the stimulus (10s). Each trial consisted of two options shown on the screen. The participants had to choose between the two options, which would give different numbers of tokens. The tokens could be exchanged for prizes at the end of the session. Thus, they choose between receiving a small number of tokens now (either 2, 4, or 6 tokens, received at the end of the scanning session) or a larger amount of tokens to be delivered in a varying delay in time (either 4, 7, 14, or 28 days). Each immediate option is paired to each delayed option 4 times. The 48 trials are presented in two rounds of 24 trials each, in a pseudo-randomised order. Each participant was shown the immediate option randomly on the right/left side of the screen (but consistently within one run) and on the opposite side on the subsequent run. Participants were instructed that immediately after the session, one trial would be chosen at random and implemented. The child received the number of tokens of the chosen option for that trial, and the subsequently chosen prize would be either taken home immediately or delivered by the post after the specified delay.

### Behaviour Data Analyses

Two of the 28 children withdrew from the study after the first task (in both cases, the intertemporal choice task). For three participants there were technical failures collecting the performance data during the Flanker task, which resulted in one participant being excluded due to the complete loss of performance data, and for 2 participants only 1 run of the task could be used in the analysis.

### Regressions on behavioural performance

Statistical analyses were conducted using RStudio (Version 1.1.442)^51^. The main performance measures for the tasks are as follows: 1) N-Back: percent of correct responses and false alarms, mean reaction times (RT) and standard deviation of RT (SDRT). These are combined to calculate a d-prime index and an intra-individual coefficient of variation for each working memory condition. 2) Flanker: percent correct responses in each condition, mean RT, and SDRT. 3) Intertemporal choice task: percent of trials in which the delayed option was chosen, mean RT and SDRT. In the Flanker and intertemporal choice tasks, the RT variables were used to calculate an intra-individual coefficient of variation for each task just as in the N-Back task.

We investigated the effects of the working memory training (WMT) on the main performance variables of the different tasks. To do so, we conducted a general linear model for the N-Back and Flanker tasks with training group (WMT vs. Non-WMT Comparison) as fixed-effects factor and task condition (N-Back: high working memory vs. low working memory; Flanker: congruent vs. incongruent) as random-effects factors. For the ICT we conducted an univariate ANOVA with percent delayed option chosen as the main performance variable and training group (as above) as between-subjects factor.

### Decision diffusion modelling

We used a Bayesian hierarchical approach to fit the parameters of the decision diffusion model (DDM) to the Flanker task using JAGS^52^ and the JAGS Wiener module^53^ together with the rjags package^54^ in R. The fitting was run with 3 chains, 100,000 burn-in samples and 10,000 posterior samples with a thinning rate of 10 samples. Drift rates were calculated as a weighted linear combination of the target and non-target stimuli in order to distinguish the relative contribution of each to the evidence accumulation rate.

### fMRI data collection and analysis

#### MRI scanning parameters

Images were acquired using a Philips Achieva 3T whole-body scanner with an eight-channel sensitivity-encoding head coil (Philips Medical Systems) at the Laboratory for Social and Neural Systems Research, University Hospital Zurich. The paradigms were written using Matlab, and presented using the Psychophysics Toolbox extension (Psychtoolbox v3.0, Brainard 1997) via a back-projection system mounted on the head-coil. We acquired gradient echo T2*-weighted echo-planar images (EPIs) with blood-oxygen-level-dependent (BOLD) contrast (37 slices per volume, Field of View 200 x 200 x 133 mm, slice thickness 3 mm, .6 mm gap, in-plane resolution 2.5*2.5 mm, matrix 80*78, repetition time 2344 ms, echo time 30 ms, flip angle 77°) and a SENSE reduction (i.e. acceleration) factor of 1.5. Volumes were acquired in axial orientation at a −20° tilt to the anterior commissure-posterior commissure line. The functional runs comprised a number of volumes in ascending order (volumes N-Back: 144, volumes FT: 145, volumes ITC:154) in addition to five “dummy” volumes at the start of each run. To measure and later correct for the homogeneity of the magnetic field B0/B1 maps were collected (short echo time = 4.29 ms, long echo time = 7.4 ms). Breathing frequency and heart rate were measured with the in-built system of the scanner in order to correct for physiological noise. A T1-weighted turbo field echo structural image was acquired for each participant (181 slices, Field of View 250 x 250 x 181 mm, slice thickness 1 mm, no gap, in-plane resolution 1*1 mm, matrix 256*256, repetition time 8.2 ms, echo time 3.8 ms, flip angle 8°).

#### fMRI preprocessing

Image analysis was performed using SPM12 (Wellcome Department of Imaging Neuroscience, Institute of Neurology, London, UK). Functional images were realigned and unwarped, segmented according to the standard T1-weighted structural images, normalized to the mean subject’s EPI template and smoothed using a 5 mm full width at half maximum isometric kernel. To account for physiological noise we used the PhysIO toolbox implementation of RETROICOR (http://www.translationalneuromodeling.org/tapas/).

#### fMRI General linear models (GLM)

We computed two GLMs at the first level (single subject) with SPM12 and results were examined at the second (group) level using the Randomise function from the FMRIB Software Library (http://fsl.fmrib.ox.ac.uk/fsl/) to implement non-parametric permutation tests (n=5000 permutations, with threshold-free cluster enhancement − TFCE). All the reported results are Family Wise Error (FWE) corrected at the voxel level and coordinates are given in Montreal Neurological Institute (MNI) space. Furthermore, for all the analyses, regressors were convolved with the canonical hemodynamic response function implemented in SPM12, high passed filtered (128s) and modelled using AR(1) autoregression.

##### Working memory task

In order to identify the potential effects of the working memory training on working memory capacities, we used a boxcar function to model BOLD activity during the low working memory (LWM) blocks (including both 0 and 1-back blocks) and separately during the high working memory (HWM) blocks (including 2 and 3-back blocks). Each block was modelled with a start with the beginning of the first stimuli of the block (that is, after the instructions screen for each block) until the end of the last stimuli of the block. Motion parameters and the physiological regressors output from RETROICOR were also included as variables of no interest. The contrasts of interests were the activation during each condition (HWM and LWM). These contrasts of interest were calculated at the individual level and the outcomes were entered into non-parametric permutation tests conducted using the randomise command in FSL at the group level for statistical inference.

##### Flanker task

Correct congruent and correct incongruent trials were modelled separately as events of duration equal to their reaction times. Furthermore, missed and incorrect trials were also included as variables of no interest as well as motion parameters and the physiological regressors output from RETROICOR. Following the GLM estimation, we computed the contrasts of interest: 1) activation during congruent, and 2) activation during incongruent trials, separately. These contrasts of interest were calculated at the individual level and the outcomes were entered into non-parametric permutation tests conducted using the randomise command in FSL at the group level for statistical inference.

### Regressions on long-term follow-up measures

In order to investigate whether intra-individual variability measures could be indicative of future outcomes at the subsequent follow-up assessments across all intervention groups, we conducted Bayesian linear regression analyses. We focused on the academic skills of arithmetic, geometry, and reading abilities, following Berger et al^10^. We also analysed the total score from the Strength and Difficulties Questionnaire (SDQ)^55^, a general mental health screening tool for indexing the level of behavioural and psychological adjustment in healthy and clinical populations. Specific follow-up or intra-individual variability measures were missing for some children (maximum missing values for any measure = 4). In order to use as much of the data as possible, we imputed the missing values using the ‘mice’ package^56^ in R. We generated ten different imputed datasets and fit Bayesian linear regressions to each of them using the R package, ‘brms’^57^ as an interface to STAN^58^. We drew our final inferences from the combined posterior distributions of all ten regression models in order to reduce the influence of any one set of imputed values on our results. The full set of regressor variables and results from these regressions are reported in Table 4.

### Conceptual replication in the ABCD study

We used the Data Exploration and Analysis Portal (https://deap.nimhda.org), which allows the fitting of additive mixed models using the R package GAMM4. We fit additive mixed models testing the relationship between the individual coefficient of variation (ICV) in the N-Back task and general mental and physical health as indexed by the Child Behavioral Checklist (CBCL)^59^, and body mass index (BMI). These models controlled for random effects for sibling pairs, fixed effects of parental race and education level, and participants’ age and sex. We selected only those participants whose performance in the N-Back task was deemed as adequate by the ABCD study’s established QA procedure (overall response accuracy for 0-back or 2-back > 60%).

## Results

### Behavioural Results

Children in the working memory training group (WMT) did not differ from those in the comparison group (CMP) at baseline. Before the start of any training program, all participants were assessed using a number of tests that included general intelligence (a modified version of the Raven Matrices), working memory (visual, spatial), inhibition (Go-NoGo task), school performance (including reading, arithmetic, geometry, etc) and general mental health screening measures (Strength and Difficulties Questionnaire). Statistical comparisons between the two groups show that they did not differ in any of these baseline measures (Table S1).

### Accuracy and response-time variability findings

Overall, adaptive working memory training led children to perform more accurately and with less trial-to-trial variability in response times during the N-Back (working memory) and Flanker (selective attention / response inhibition) tasks. Figure 1 shows the differences between training groups in the N-Back, and Flanker tasks (see Table S2 for the full set of descriptive statistics and results). The WMT group responded more accurately in the Flanker task across both the congruent (i.e. easier) and incongruent (i.e. harder) trials. In the N-Back task, children in the WMT group were more accurate than those in the non-working memory comparison group on low working memory trials (0-1 back), but the two groups did not significantly differ on high working memory trials (2-3 back).

**Figure 1.**
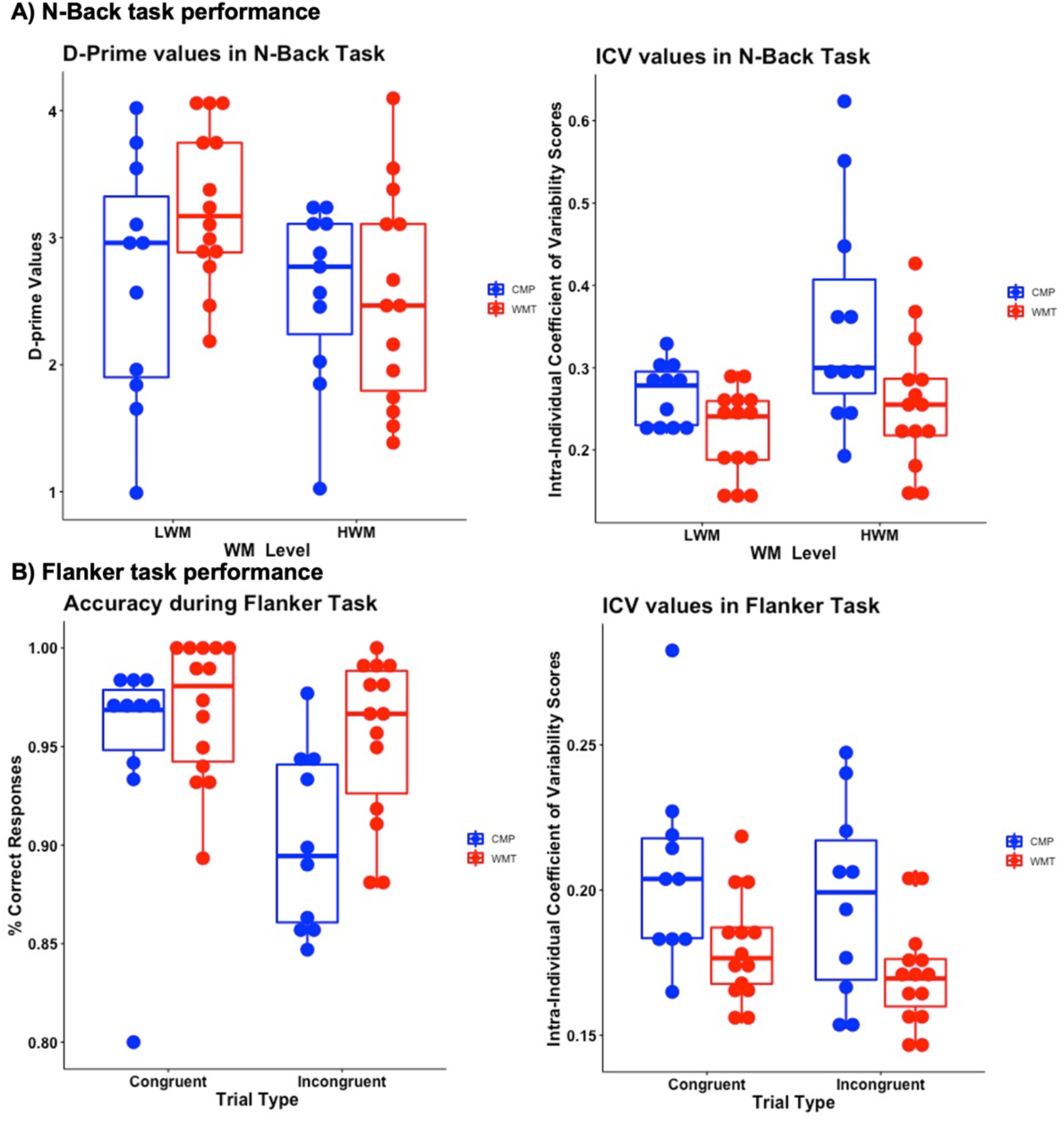
Task performance in the N-Back and Flanker tasks. The figure shows the performance for each group in the main outcome variables of the N-Back and Flanker tasks. Top left panel: The working memory training group (red) shows higher D-prime scores in the low working memory condition than the comparison group (blue), while both perform similarly in the high working memory condition. Top right panel: Children in the working memory group show reduced intra-individual coefficient of variation relative to those in the comparison group on both task conditions. Bottom left panel children in the working memory group show higher accuracy than those in the comparison group in the Flanker task. Bottom right panel: children in the working memory group show reduced intra-individual coefficient of variation than those in the comparison group across all trial types in the Flanker task.

In addition to better accuracy, children that received adaptive working memory training also showed less intra-individual variability in response times than children that did not receive working memory training (Figure 1). We computed the intra-individual coefficient of variation as intra-individual RT standard deviation / intra-individual RT mean. Note that the standard deviation of reaction times was not significantly correlated with mean response times (N-Back: r = −0.2816, p = 0.1727, 95% CI [-0.61, 0.13]; FT: r = −0.01, p = 0.959, 95% CI [-0.41, 0.39]). Furthermore, an analysis of ex-gaussian parameters fit to each child’s response time distribution revealed that the standard deviation (sigma) and exponential (tau) parameters differed between the working memory trained and comparison groups, but there was no significant difference in the means (mu) of the response time distributions (Table S3). In other words, more variable individuals were not reliably faster or slower to respond overall. Rather they were more inconsistent in the way they executed their responses. Intra-individual variability was highly correlated across the N-Back and Flanker tasks (r = 0.65, p = 0.0008, 95% CI [0.32, 0.84]). Together, the pattern of results across both cognitive tasks suggests that the adaptive working memory training intervention may have improved children’s ability to engage and maintain attention on task-relevant information in a domain-general manner soon after the five weeks of training were complete.

We did not detect any effects of adaptive working memory training on intertemporal monetary choice outcomes or on intra-individual variability in that decision task. Thus, children in the WMT group did not differ from those in the comparison group in their willingness to wait for larger, delayed monetary rewards within the range of amounts and delays included in this study. That being the case, we focus on the N-Back and Flanker tasks in the remaining sections of this paper.

### fMRI results

Along with better accuracy, the WMT group showed increased activation compared to the comparison group in brain regions that are part of attention and control networks during the low working memory trials. This included portions of fronto-striatal-thalamic systems such as the right caudate, putamen, pallidum, thalamus, inferior middle and superior frontal gyri, the dorsal anterior cingulate and the supplementary motor cortex (Table 1, Figure 2). Consistent with the behavioural findings of similar accuracy in the high working memory condition, there were no significant differences in the BOLD signal across groups during the high working memory trials. We did not detect any significant differences in activity as a function of working memory training during the Flanker task.

**Table 1.** Regions showing increased BOLD signal response in the WMT versus CMP groups during the low working memory condition of the N-Back task.

| Region | Cluster Size | x; y; z | T-stat (1) | T-stat (2) |
| --- | --- | --- | --- | --- |
| Juxtapositional Lobule Cortex (formerly Supplementary Motor Cortex); Precentral Gyrus; Superior Frontal Gyrus; Cingulate Gyrus, anterior division | 189 | - 10; -8; 58 | 7.1 | 6.01 |
| Frontal Pole; Middle Frontal Gyrus; Superior Frontal Gyrus | 146 | 37; 36; 43 | 4.66 | 4.57 |
| Right Putamen; Right Caudate | 133 | 15; 9; 18 | 4.79 | 4.18 |
| Right Pallidum; Right Accumbens; Right Putamen; Right Hippocampus; Right Amygdala | 125 | 10; -13; -12 | 5.91 | 5.42 |
| Superior Frontal Gyrus; Paracingulate Gyrus; Frontal Pole | 77 | -2; 41; 45 | 4.8 | 4.06 |
| Precentral Gyrus; Inferior Frontal Gyrus, pars opercularis; Postcentral Gyrus; Central Opercular Cortex | 46 | 47; 4; 18 | 4.78 | 3.72 |
| Right Putamen | 24 | 25; -8; 23 | 5.02 | 5.01 |
| Cingulate Gyrus, anterior division; Paracingulate Gyrus; Juxtapositional Lobule Cortex (formerly Supplementary Motor Cortex) | 23 | 12; 11; 35 | 4.64 | 4.34 |
| Inferior Frontal Gyrus, pars opercularis; Inferior Frontal Gyrus, pars triangularis; Precentral Gyrus | 16 | 52; 19; 10 | 4.29 | 3.11 |
| Precentral Gyrus; Postcentral Gyrus | 12 | -25; -26; 50 | 5.15 | 4.04 |
| Middle Frontal Gyrus; Inferior Frontal Gyrus, pars triangularis; Inferior Frontal Gyrus, pars opercularis | 7 | 42; 29; 23 | 4.42 | 3.56 |
| Precentral Gyrus | 6 | 22; -11; 45 | 5.26 | 4.11 |

All reported regions are significant at $p < 0.05$ after whole brain FWE correction at the voxel level. Here we report only regions with a cluster extent greater than 5 voxels. The full sets of unthresholded t-statistics can be found on Neurovault.org (https://identifiers.org/neurovault.collection:9005). The FWE correction was based on 5000 permutations of the threshold free cluster enhancement (TFCE) values. The TFCE values and permutation-derived test statistics were calculated using the Randomise function implemented in FSL. All coordinates are listed in MNI space and represent the peaks of all clusters formed by contiguous voxels. Anatomical labels were derived from the Harvard–Oxford cortical and subcortical atlases. The two T-stat columns show the T value for the comparisons (1) in the whole sample (working memory training group = 14, comparison group = 11, total N= 25); (2) in the sample after excluding the three children with non-random assignment to the pure control condition (working memory training group = 14, comparison group = 8, total N= 22)

**Figure 2.**
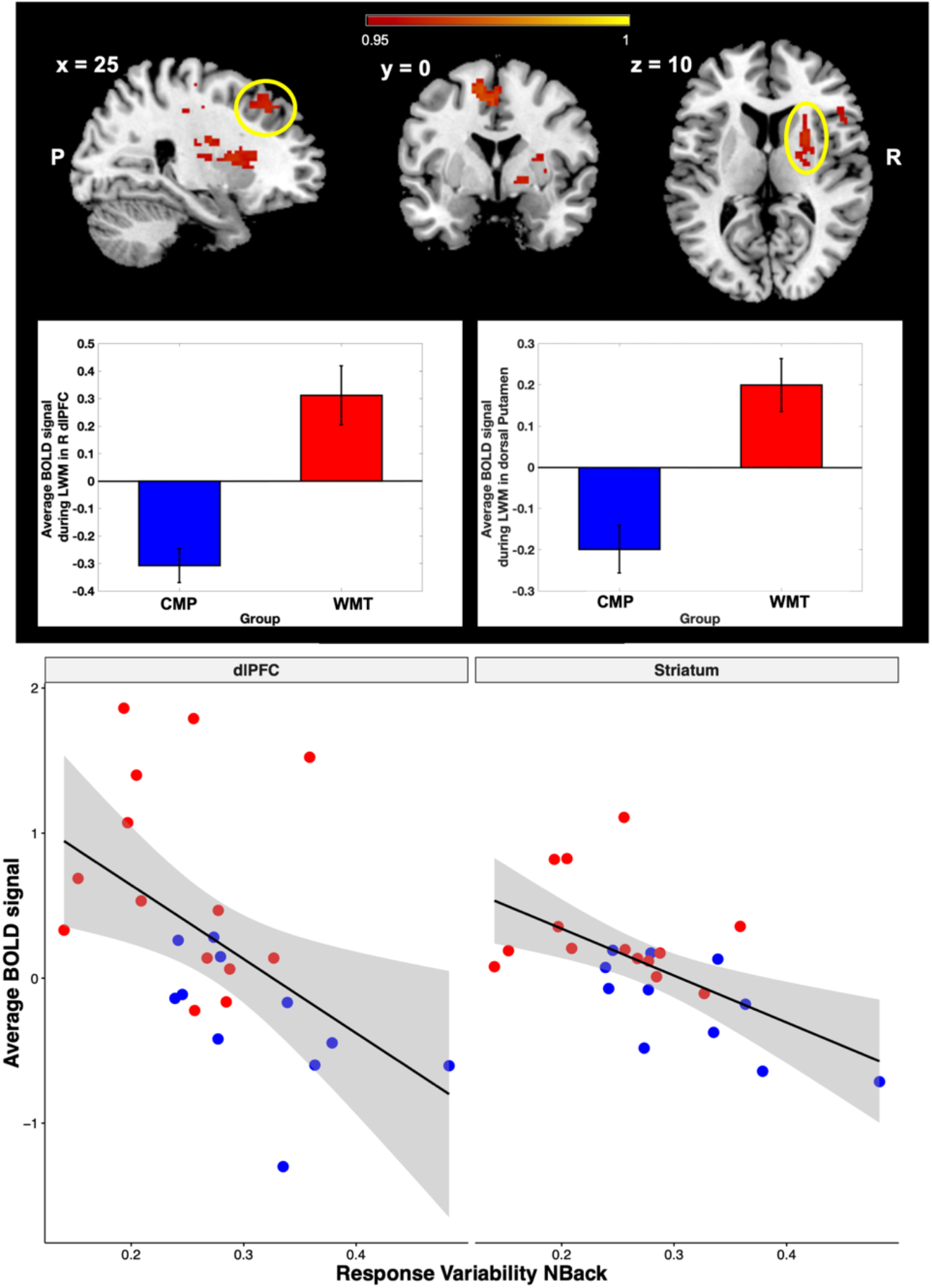
Group differences on brain activation. The WMT group had greater activity in frontostrial regions than the CMP group in N-Back tasks. Consistent with the behavioural results, these differences were specific to the low working memory condition. The bar graphs show the average BOLD signal in each group, in the two clusters circled in yellow (1) right dorsolateral prefrontal cortex (left panel) and (2) right putamen (right panel). The scatter plots in the bottom row show the association between BOLD signal and Individual differences in the coefficient of variation across all trials. Children in the WMT group are shown in red while those in the CMP group are shown in blue.

In addition, we found that task-related BOLD signal levels in regions that showed greater activity in the WMT group (see Figure S2) also correlated with the intra-individual coefficients of variation and/or accuracy on the N-back task across all participants (Figure 2, bottom row). Some relation to accuracy and the intra-individual coefficient of variation in these regions is to be expected given that there are group differences in intra-individual variability (see Figure 1). However, activity in the dorsal striatal functional ROI, encompassing dorsal caudate and putamen, was significantly associated with intra-individual variability even after accounting for the effects of working memory training condition (coef = −0.25, p = 0.004; Table 2). There were similar, though not significant, trends in the dorsolateral prefrontal cortex (dlPFC) for intra-individual variability, and in the anterior cingulate cortex/supplementary motor area for accuracy (Table 2).

**Table 2.** Associations between BOLD signal in ROIs where group differences were identified in the LWM contrasts and intra-individual coefficient of variation and d-prime across all trials.

| **Dorsal striatum** |  |  |  |
| --- | --- | --- | --- |
|  | Estimate (se) | T value | P value |
| Group | 0.17 (0.15) | 1.16 | - |
| ICV | -0.25 (0.08) | -3.26 | 0.004* |
| d-prime | 0.12 (0.06) | 2.10 | 0.049* |
| **dIPFC** |  |  |  |
|  | Estimate (se) | T value | P value |
| Group | 0.56 (0.31) | 1.782 | - |
| ICV | -0.32 (0.16) | -1.978 | 0.0619 |
| d-prime | 0.09 (0.13) | 0.689 | 0.4989 |
| **dACC-SMA** |  |  |  |
|  | Estimate (se) | T value | P value |
| Group | 0.50 (0.20) | 2.497 | - |
| ICV | -0.13 (0.10) | -1.249 | 0.2260 |
| d-prime | 0.14 (0.08) | 1.794 | 0.0879 |

The table reports the results from three linear regression models testing the association between the intra-individual coefficient of variation and d-prime during the N-Back task and BOLD signal after accounting for the effects of training condition (Group). The linear model in each region was computed as BOLD signal ∼ Group + ICV + d-prime + Delay, where Delay is a control variable that accounts for the delay between the end of the intervention and the scanning session (omitted from the table for conciseness). We list the T-values for the binary regressor indicating training group for comparison with the linear regressor for ICV. However, we do not report p-values for the Group regressor because the functional ROIs were originally identified with this contrast, making the analysis circular. The point of these regressions is to test if differences in ICV provide additional explanatory power in the region where activity differed between training groups. se = standard error; ICV = Individual Coefficient of Variation; dIPFC = dorsolateral prefrontal cortex; dACC= dorsal anterior cingulate; SMA = supplementary motor area. \* denotes significance at 0.05 p-value threshold.

### Diffusion Decision Model analyses

Diffusion decision models are a form of computational modelling that can quantify and distinguish between different cognitive processes that may give rise to intra-individual variability^60,61^. Previous work has shown that the effects of attention on task performance and decision making can be formalized and quantified by diffusion decision models^62-64^ (DDM). Here, we fit the DDM to children’s behaviour in the Flanker task. We did not fit data from the N-Back task because it required responses only on target trials, which were a small minority (25%) of all trials.

The idea behind sequential sampling models is that participants repeatedly extract noisy signals from the stimuli and/or their own minds until they are sufficiently sure of the best response. Previous work has shown that visual attention changes the way in which the evidence for each alternative is accumulated^62-64^. From moment to moment, items that are fixated influence the accumulation rate more strongly than items that are not currently fixated. We hypothesized that, if working memory training influenced the ability or motivation to selectively attend to task relevant information, then we should see differences in the estimated drift rates for the DDM between the WMT and CMP groups. Past work examining the role of attention on DDM parameters has used visual fixation (measured with eye tracking) as a proxy for attention^62,64^, and estimated how drift rates change within a choice as a function of the stimulus the participants fixate at a given point in time. In contrast, we don’t need to subdivide the trial durations based on fixation locations in order to test our hypothesis about drift rates in the Flanker task. The design of the Flanker task is such that only one of the stimuli on the screen contains the trial-specific information that is relevant for behaviour, the target. Therefore, we can estimate how much target versus distractor stimuli contribute to the drift rate across the entire duration of the trial in each training group.

In our specification of the DDM, the magnitude of the drift rate coefficients informs us about how strongly each stimulus influences the evidence accumulation processes. In the flanker task, children should be focused on the target fish because it alone provides evidence for the correct response in each trial. The direction the flanking distractor fish are facing is irrelevant and should be ignored. Thus, we specified the drift rate according to equation (1) below. We hypothesized that β_1_ – β_2_ (i.e. the weight on relevant minus irrelevant information) should be greater in the WMT than the CMP group.

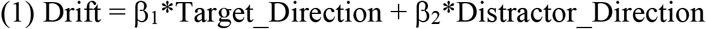

The DDM results are consistent with an improved ability to focus attention on task relevant features following working memory training. The results from the Flanker task showed that children in the WMT group were more sensitive to the information carried by the target fish (i.e. its direction) relative to distractor fish (p = 0.008, Table 3) and utilized a higher response threshold (p = 0.048, Table 3) than children in the comparison group.

**Table 3.** Diffusion Decision Model parameters for the Flanker task.

| *DDM Parameter* | **WMT** |  |  | **CMP** |  |  | **WMT - CMP** |  |  |
| --- | --- | --- | --- | --- | --- | --- | --- | --- | --- |
|  | *m* | *HDI* |  | *m* | *HDI* |  | *m* | *HDI* |  |
| Target drift coef. | 2.84 | 2.30 | 3.38 | 1.99 | 1.65 | 2.37 | **0.84\*** | 0.20 | 1.49 |
| Distractor drift coef. | 0.20 | 0.03 | 0.38 | 0.25 | 0.00 | 0.51 | -0.05 | -0.36 | 0.25 |
| Target - Distractor | 2.64 | 2.06 | 3.20 | 1.75 | 1.31 | 2.19 | **0.89\*** | 0.14 | 1.57 |
| Boundary | 2.53 | 2.13 | 2.96 | 2.09 | 1.75 | 2.42 | **0.45\*** | - 0.08 | 0.98 |
| Non-Decision Time | 0.30 | 0.22 | 0.37 | 0.24 | 0.14 | 0.35 | 0.05 | - 0.07 | 0.18 |

This table lists the mean (*m*) as well as the lower and upper bounds of the 95% highest density interval (HDI) of the posterior distributions for parameters or parameter differences from the decision diffusion model (DDM) fit to the Flanker task. The DDM was fit to the Flanker task data separately for the children in the group that received working memory training (WMT) and those in the comparison group (CMP). The asterisks next to the mean differences between WMT and CMP denote those means that are significantly different based on a one-sided test of posterior probability of the mean for WMT being greater than the CMP group.

**Table 4.**
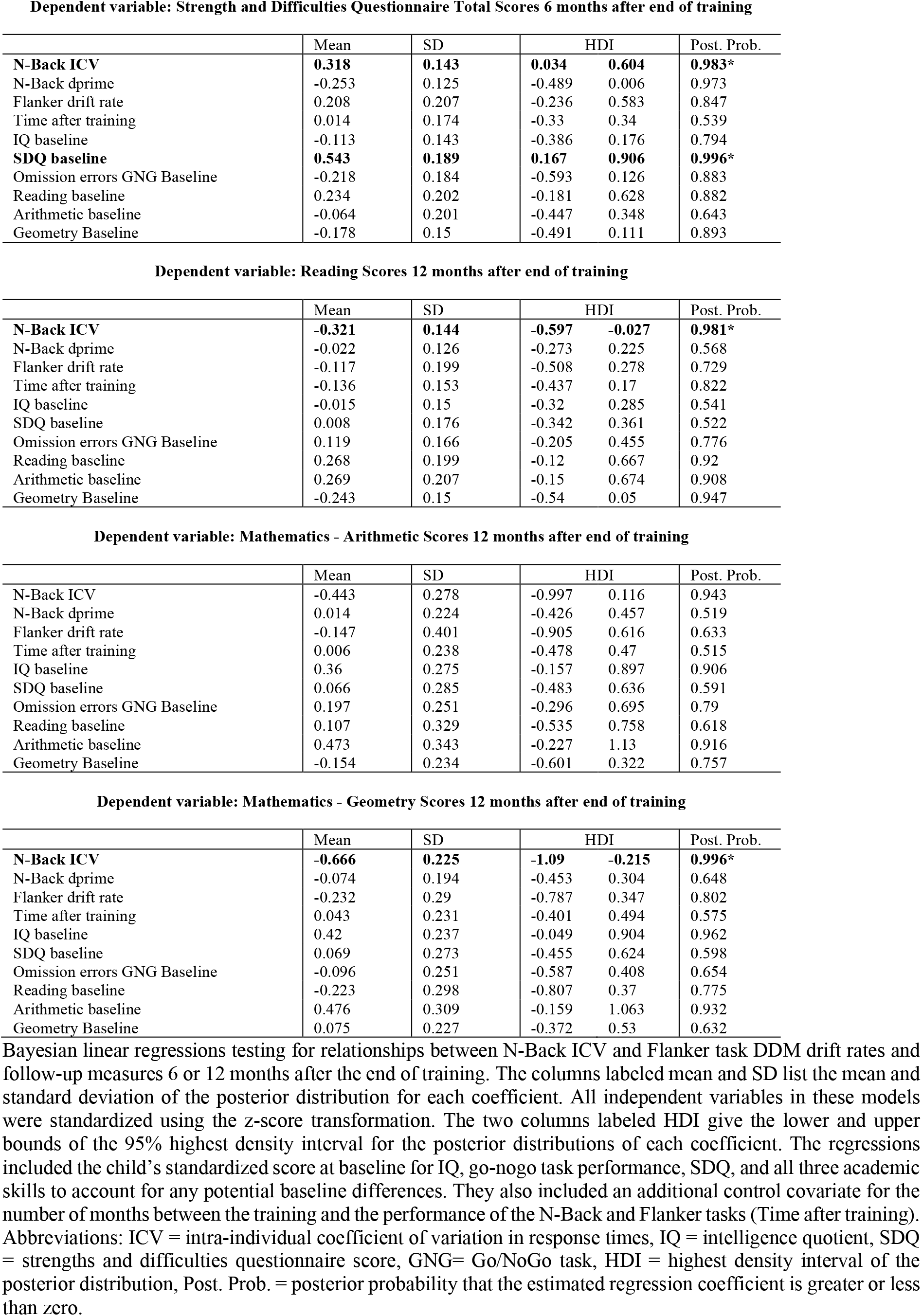
Results of the regression analyses showing the association between task performance and follow-up measures in our sample.

Furthermore, simulated responses from the fitted diffusion decision model reproduce the patterns of response time variability observed in the Flanker task. To generate simulated responses in the Flanker task, we used each participant’s best-fitting DDM parameters. We then compared the simulated response times across groups and found that the RTs were less variable for simulated agents using parameters from the WMT participants than for simulated agents based on CMP children’s parameters (Table S4).

#### Associations between post-training cognitive task performance and follow-up measures

The recent work of Berger and colleagues^10^ showed that some effects of working memory training in children are only evident in follow-up assessments 6 or 12-13 months after training was completed. Therefore, we investigated if the intra-individual variability measures that we computed immediately after the training could serve as indicators of relevant future outcomes at the subsequent follow-up assessments.

The ultimate goal of any training is to have a meaningful impact in the life of the participants. Therefore, we focused on outcome measures of academic performance and behavioural and psychological adjustment that could represent an impact in “real-life”. Specifically, we examined the total score in the Strength and Difficulties Questionnaire (SDQ)^55^, a behaviour and mental health screening measure typically administered in clinical settings to identify potential problematic areas in a child that merit further assessment by a specialist. The SDQ was filled out by parents 6 months after training. We also examined tests of academic performance in reading and two mathematics subscales (geometry and arithmetic) conducted 1 year after training. We focused on these specific academic skills because of the results from the independent sample in Berger et al^10^, which show that working memory training improved geometry and reading scores, but not arithmetic.

We conducted linear regression analyses that tested if these measures of future general mental health and academic skill could be explained by children’s accuracy (d-prime) or response time variability in cognitive tasks performed at the end of the training period. Specifically, we used the coefficient of variation in response times and d-prime scores from the N-Back task and the estimates of DDM drift rates from the flanker task to explain future outcomes. Response time variability explained significant additional variance in future SDQ scores (standardized coef. = 0.32 ± 0.14), geometry (standardized coef. = −0.66 ± 0.23), and reading (standardized coef. = −0.32 ± 0.14), even after accounting for baseline scores in those measures and IQ (Table 4). In contrast, no post-training cognitive task performance or baseline measures were significantly associated with future arithmetic scores. Thus, in our sample, the intra-individual variability in response times measured right after the intervention correlated with future performance in the same academic domains that Berger and colleagues previously found to be improved one year after working memory training in an independent sample.

#### Testing the association between intra-individual variability and general mental health in an independent sample

We examined whether there were similar associations between intra-individual variability in response time during the N-Back task and mental and physical health in the children in the ongoing longitudinal ABCD study (https://abcdstudy.org, data release version 2.0.1; N = 7894 children). In this case, we had to limit our analysis to concurrent baseline measures because follow-up measures were not yet available. The ABCD study is a longitudinal, multicentre study of children’s cognitive and neurobiological development starting from age 10. This study includes a wide range of standardized questionnaires and interviews covering both general well-being and clinical measures. In addition, participants perform several cognitive tasks, including an N-back task^65^. We tested if intra-individual variability, once again quantified as the coefficient of variation in response times, during the N-Back task was associated with scores on the Child Behavioural Checklist (CBCL)^59^. The CBCL is a measure of current behavioural and psychopathological symptoms with high correspondence to the SDQ when both scales are applied to the same individuals. The ABCD study includes the CBCL, but not SDQ. We also tested for potential relationships between intra-individual variability and body mass index (BMI) scores. We chose BMI as an additional translational measure robustly associated with physical, cognitive, and socioeconomic well-being. Decreased intra-individual variability in response times during the N-Back task was associated with decreased concurrent total CBCL scores (i.e. lower variability was associated fewer potential problems, standardized coef. = 0.27 ± 0.13). Less intra-individual variability during the N-Back was also associated with lower BMI scores (standardized coef. = 0.15 ± 0.04; Table 5). Thus, the results from this large independent sample are consistent with our original findings showing that less intra-individual variability in response time is associated with increased general mental health and academic skills.

**Table 5.** Results of the regression analyses showing the association between Intra-individual variability and general mental and physical health in the ABCD Study.

| **A** Dependent variable: CBCL total score |  |  |  |  |
| --- | --- | --- | --- | --- |
|  | Estimate | Std Error | T value | p-value |
| N-Back ICV | 0.27035 | 0.12445 | 2.17 | 0.02985 |
| N-Back correct rate | -1.07062 | 0.12994 | -8.24 | <1e-6 |
| Controlling for: |  |  |  |  |
| Race |  |  |  |  |
| Sex |  |  |  |  |
| Parental education |  |  |  |  |
| Age |  |  |  |  |

| **B** Dependent variable: Body Mass Index (BMI) |  |  |  |  |
| --- | --- | --- | --- | --- |
|  | Estimate | Std Error | T value | p-value |
| N-Back ICV | 0.1463 | 0.04249 | 3.44 | 0.000578 |
| N-Back correct rate | -0.0487 | 0.04434 | -1.10 | 0.27215 |
| Controlling for: |  |  |  |  |
| Race |  |  |  |  |
| Sex |  |  |  |  |
| Parental education |  |  |  |  |
| Age |  |  |  |  |

## Discussion

The present study examined how the neurocognitive mechanisms underlying the short-term impact of adaptive working memory training in primary school children relate to training benefits that emerge months or years after training. Overall, our results suggest that in addition to working memory itself, there may be concurrent improvements in selective and sustained attention during or directly after five weeks of training. We show that intra-individual variability in response times during working memory and selective attention tasks can be used to detect short-term training effects in children, and that such measures may be indicative of the persistence and/or emergence of far-transfer benefits months to years after training is completed.

Our findings indicate that improvements in attention are among the immediate results of adaptive working memory training. Working memory and attention processes are thought to be closely linked and interdependent^6,17,66-71^. Although they have different primary targets, both the Flanker and N-Back tasks require the ability to maintain attentional focus throughout the duration of the task (sustained attention), and to identify the target stimuli and filter out or inhibit responses to non-target stimuli (selective attention). At the neural level, differences between the WMT and CMP groups were found in striatum as well as the lateral and medial prefrontal cortices, which are brain regions that, among other things, support selective and sustained attention functions^18,19,72^. These neural differences were accompanied by better signal detection performance (i.e. higher d-prime), reduced intra-individual variability in response times, and more efficient accumulation of relevant information (i.e. higher DDM drift rates) in children that received adaptive working memory training. All of these behavioural measures are related to and dependent on attention. Therefore, taken together, our neural and behavioural results suggest that the benefits of the working memory training program used in this study are at least partially mediated by improvements in attention processes leading to consistent and effective responses to task relevant information and reduced processing of irrelevant, distracting stimuli.

These results lend further support to theories of the mechanisms underlying training benefits. A meta-analysis of previous training studies concluded that the Cogmed-RM adaptive working memory training program has effects on attention in daily life^73^. The improvements in attention processes we detected at the end of the training are consistent with previous results and theories about the basis of far transfer effects following cognitive training as well^32,33,74^. Specifically, these far-transfer benefits occur when the trained and transfer skills share common fundamental cognitive processes. Given the important role of attention as a prerequisite to many cognitive processes, it could serve as a basis for far-transfer effects following working memory training.

Recently, the effects of the adaptive working memory training in school-age children have been shown to emerge over 6 to 12 months^10^. Initial improvements in attention may serve as a scaffold for later changes in higher cognitive processes that facilitate better school performance. Our current results suggest that attention functions might be among the first to improve from this type of training, and that later emerging benefits to academic skills and general mental health are associated with immediate improvements in attention processes. It is not surprising that working memory training would also influence attention control (e.g. selective attention, sustained attention, or goal-directed attention reallocation) given that these processes are postulated to be pre-requisites for the successful implementation of working memory^6,17,66-68,71^. There is also evidence that the associations between working memory capacity and various cognitive and academic skills are partially mediated by a common reliance on attention control^66,70,75^. Given the apparent role of attention processes in mediating the far transfer of training effects, it is important to measure these processes when assessing the efficacy of working memory training and other forms of cognitive training.

The ability of intra-individual variability metrics to detect individual differences in attention control could explain the association we find between them and the future emergence of benefits to academic skills and general mental health after working memory training. Intra-individual response time variability metrics are sensitive and reliable measures of individual differences in attention control processes^34,35^. They are often used as an index of an individual’s attention allocation efficiency or degree of fluctuation in attention control during task performance^45,46,76-78^. Intra-individual variability has been linked with cognitive control measures in healthy children and adults, and the variability in response times measured in one task is correlated with working and long-term memory or intelligence measured in separate tasks^28,42,45,46,79-81^. It also differs between healthy individuals and those with attention deficit hyperactivity disorder (ADHD)^36,37,42,82,83^. However, increased response time variability is not unique to ADHD and is seen in various psychiatric and neurological disorders (e.g. traumatic brain injury, dementia, and schizophrenia), in which attention deficits may play an important, though less prominent, role^36,38,44,83,84^. Increased intra-individual variability is commonly observed in non-affected relatives as well as patients, indicating that it may capture shared genetic or environmental risk factors for current and future psychopathologies^44,76,82,85,86^. In fact, a recent review by Haynes et al. highlights several longitudinal studies in older adults that have shown that the intra-individual variability in response times is associated with future levels of cognitive impairment and mortality^84^. Thus, intra-individual variability measures are sensitive to not only to current cognitive and neurological function, but also associated with the future stability or decline in those functions.

Here, we have shown that intra-individual variability metrics can detect the short-term efficacy and are indicative of the emergence of longer-term benefits of working memory interventions aimed at improving cognitive skills and academic performance in children. Five weeks of working memory training led to significant decreases in intra-individual response time variability on two separate cognitive tasks (N-Back and Flanker), completed soon after the training period ended. Consistent with their ability to forecast cognitive decline in the elderly, we found that measures of the intra-individual variability computed at the end of training were associated with improvements in academic skills and general mental health in children up to one year after training. Lower post-training variability was related to better future scores on tests of academic skills and strengths/weaknesses in classroom and social behaviour for the full sample.

Our results suggest that measures of intra-individual variability are useful in evaluating intervention efficacy. However, there are several important questions that still need to be addressed. For example, can we use intra-individual variability metrics to determine when an individual has received a sufficient dose of the training intervention? If so, then we could tailor the amount of training to each person in order to improve the cost benefit trade-offs inherent in any training program. Another key question our findings raise is what types of tasks (e.g. those targeting working memory, attention, task-switching, etc) and measures of intra-individual variability are best suited to assessing the short and long-term outcomes of cognitive training. Previous work has quantified intra-individual variability in response times in several different ways^82,83,87^. We found significant differences in response time variability between training groups in both selective attention (Flanker) and working memory (N-Back) tasks, that were robust across common measures of variability (coefficient of variation, ex-gaussian decomposition, and diffusion decision modelling). However, there may be differences in how well the different measures of variability and/or task designs predict the emergence of benefits to specific areas of academic performance or general mental health in the longer term. This question will be important to address in future studies that collect and compute multiple longitudinal measures in large samples of participants.

Although a relative strength of our study is the amount of longitudinal data we have on each participant, one of the limitations is the small sample size. The concerns that small sample sizes might raise are mitigated, in this case, by the fact that key results replicate in much larger samples. Specifically, the working memory training effects we find replicate those found in Berger et al^10^ using the same form and duration of training in a separate sample of over 500 children. We also conceptually replicate the associations between the coefficient of variation in response times during an N-Back task and measures of mental and physical health in over 7800 children from the ABCD study^65^. Another limitation of this study is that, although it contains a number of pre-training baseline measures, it lacks data from the Flanker and N-Back tasks before training. The lack of baseline data on these tasks prohibits us from testing if the level intra-individual variability before the intervention is related to either short or long-term training outcomes. It will be important to determine if baseline measures of intra-individual variability can be used to help assign individuals to the appropriate level of initial training duration or potentially even training types. Moreover, changes in variability (i.e. post minus pre-training) may be even better predictors of the emergence of future benefits than post training measures alone^10,88^.

Effective means of enhancing cognitive abilities have been a long-standing goal in many disciplines. Our current work adds to the existing evidence that adaptive working memory training can significantly benefit school-aged children^6,10,89,90^. Moreover, it provides additional insights into the mechanisms underlying these benefits. Together with the recent findings of Berger et al^10^, it also highlights the importance of including long-term follow-ups in any evaluation of training efficacy. In addition to long-term follow-up data, we demonstrate the utility of using response time variability metrics as an immediate indicator of intervention success. The practical relevance of such an immediate assessment tool should not be overlooked, as it could potentially allow for tailoring training interventions in terms of duration or content without needing to wait for years for follow-up data to determine whether or not long-term benefits will emerge.

## Author contributions

AC and TH designed the task-based fMRI study and analysis plan with input from HH. AC collected the combined fMRI plus Flanker, Intertemporal choice, and N-Back task data. EB, HH, EF, DS and KW designed and implemented the in-school training interventions and follow-up measures. AC and TH analyzed the data. AC and TH wrote the manuscript with input from EB, HH, EF, DS and KW.

## Acknowledgements

This study was supported by the National Center of Competence in Research Affective sciences hosted by the University of Geneva (SNSF grant number 51NF40-104897).

## Data Availability

Code for the data analysis is openly available at https://osf.io/35f7g/

## Supplementary Material

**Figure S1.**
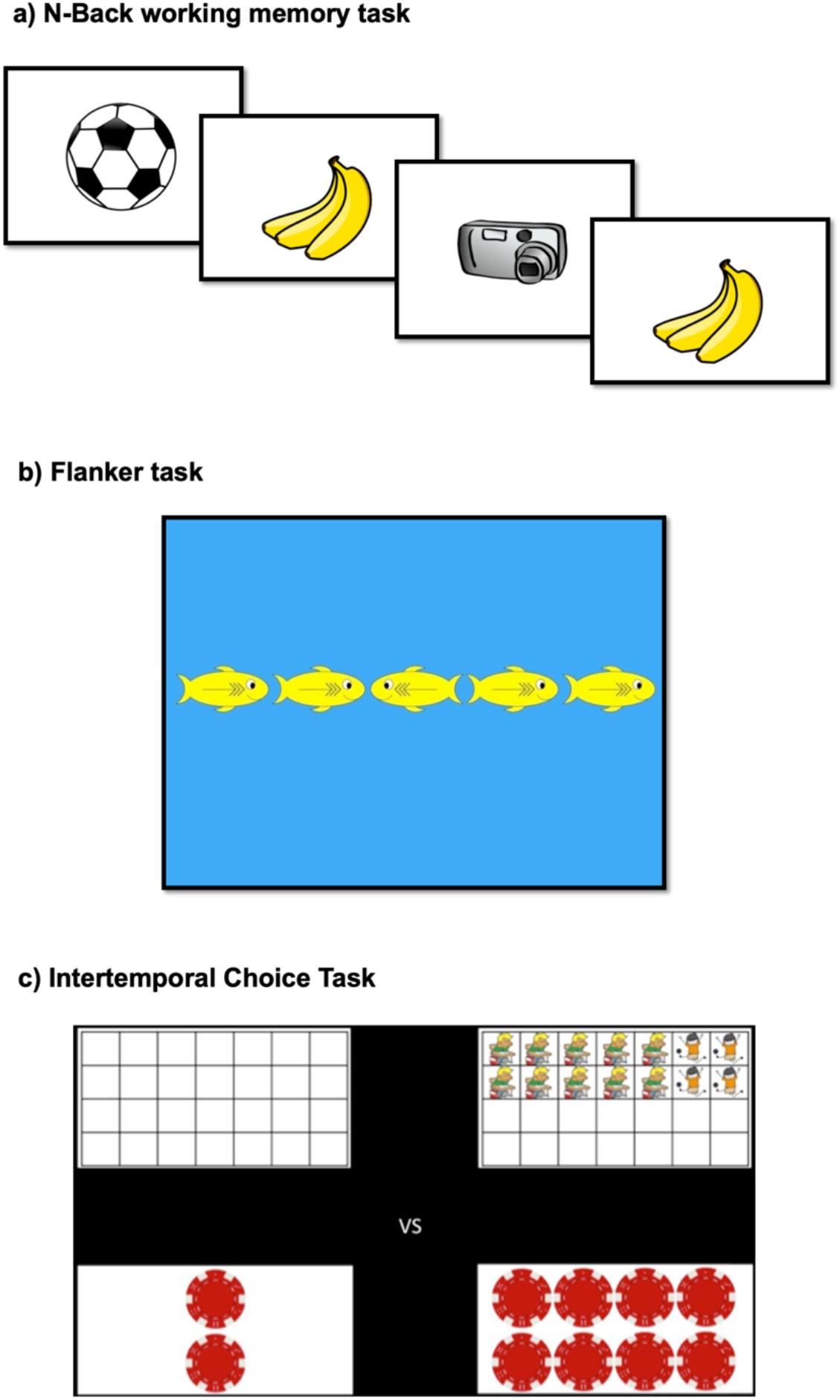
Schematic representation of the fMRI tasks. **Schematic representation of the fMRI tasks**. (a) N-Back working memory task. The image respresents a trial in the 2-Back condition, where the participant has to press the button when the image presented is the same as 2-before it. (b) Flank task. The image represents an incongruent trial, where the direction of the central fish (target stimuli) is opposed to that of the flanker stimuli, (c)lntertemporal choice task. The image respresents a trial where the participant can receive either 2 tokens immediately (shown on the left side of the screen), or 8 tokens after a 14-days delay (shown on the right side of the screen).

**Figure S2.**
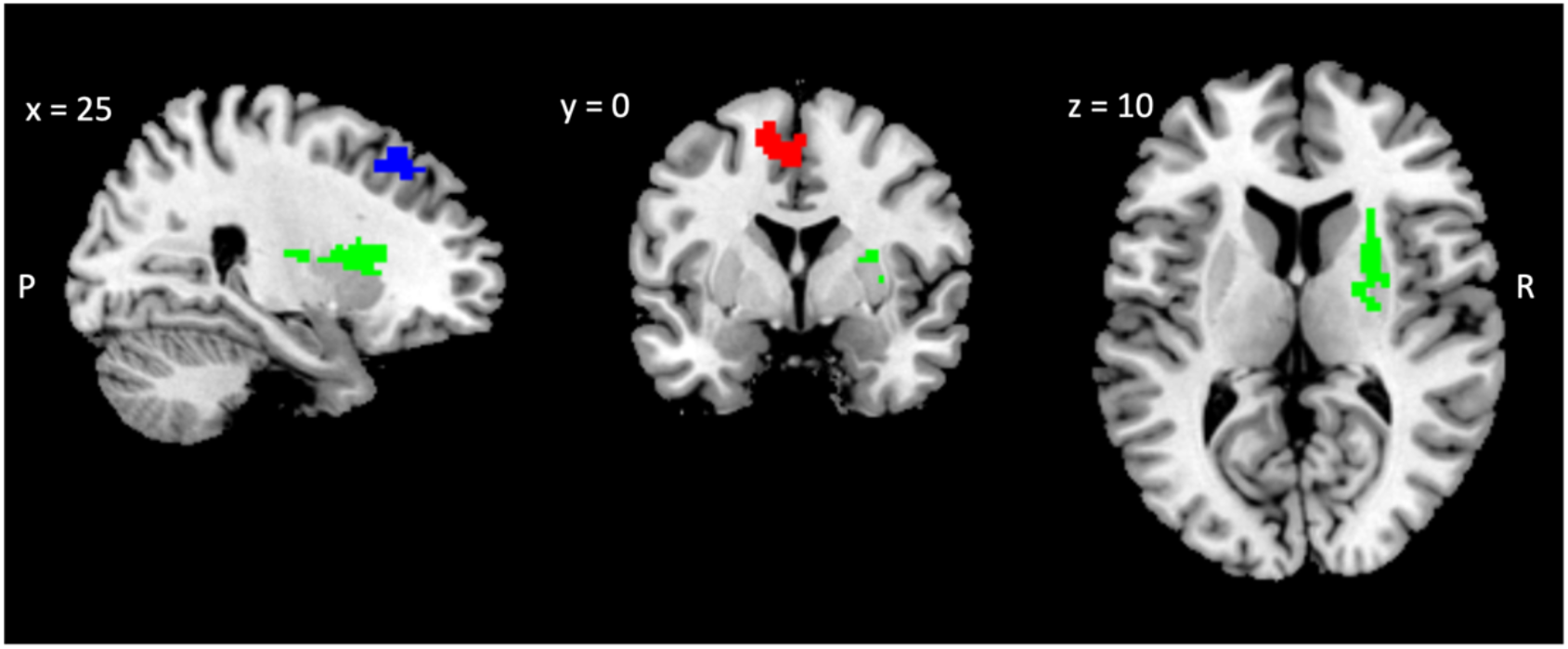
ROIs included in the analysis of the association between individual differences in task activation and coefficient of variation. we computed the average BOLD response across significant voxels in the dorsal striatum (green), the dorsolateral prefrontal cortex (blue) and the dorsal anterior cingulate /supplementary motor area (red). These were regions of interest (ROIs) were selected from the contrast between the WMT and CMP groups on low-working memory trials.

## Supplemental Tables

**Table S1.** Scores and group differences on school-based performance and mental health measures at baseline.

| Task | Group | Mean (std) | F value (df) | P value |
| --- | --- | --- | --- | --- |
| Ravens | CMP | -0.11 (1.10) | 0.603 (2,25) | 0.5549 |
|  | WMT | 0.22 (0.96) |  |  |
| Odd-one-out | CMP | 0.39 (0.78) | 0.777 (2,25) | 0.4707 |
|  | WMT | -0.04 (0.99) |  |  |
| Digit Span | CMP | 0.43 (1.15) | 0.0536 (2,25) | 0.9479 |
|  | WMT | 0.39 (1.01) |  |  |
| Lottery game | CMP | 0.13 (0.99) | 0.2229 (2,25) | 0.8018 |
|  | WMT | 0.12 (1.17) |  |  |
| Saving box game | CMP | 0.19 (1.05) | 0.0234 (2,25) | 0.9769 |
|  | WMT | 0.11 (1.12) |  |  |
| Commission errors GNG | CMP | -0.06 (1.30) | 0.4481 (2,25) | 0.6439 |
|  | WMT | 0.02 (0.98) |  |  |
| Omission errors GNG | CMP | -0.09 (1.10) | 1.801 (2,25) | 0.186 |
|  | WMT | -0.12 (0.76) |  |  |
| SDQ total score | CMP (N=12) | 0.07 (0.89) | 0.7868 (2,21) | 0.4682 |
|  | WMT (N=12) | -0.34 (0.68) |  |  |
| SDQ internalizing score | CMP (N=12) | -0.30 (0.71) | 2.246 (2,22) | 0.1296 |
|  | WMT (N=13) | -0.59 (0.45) |  |  |
| SDQ externalizing score | CMP (N=12) | 0.35 (1.08) | 1.202 (2, 21) | 0.3204 |
|  | WMT (N=12) | -0.12 (0.94) |  |  |
| Reading | CMP | 0.24 (1.09) | 0.186 (2,25) | 0.8313 |
|  | WMT | 0.46 (0.95) |  |  |
| Math Arithmetic | CMP | 0.46 (0.85) | 0.0029 (2,25) | 0.9971 |
|  | WMT | 0.49 (0.95) |  |  |
| Math Geometry | CMP | 0.12 (0.93) | 1.542 (2,25) | 0.2337 |
|  | WMT | 0.33 (0.96) |  |  |

**Table S2.**
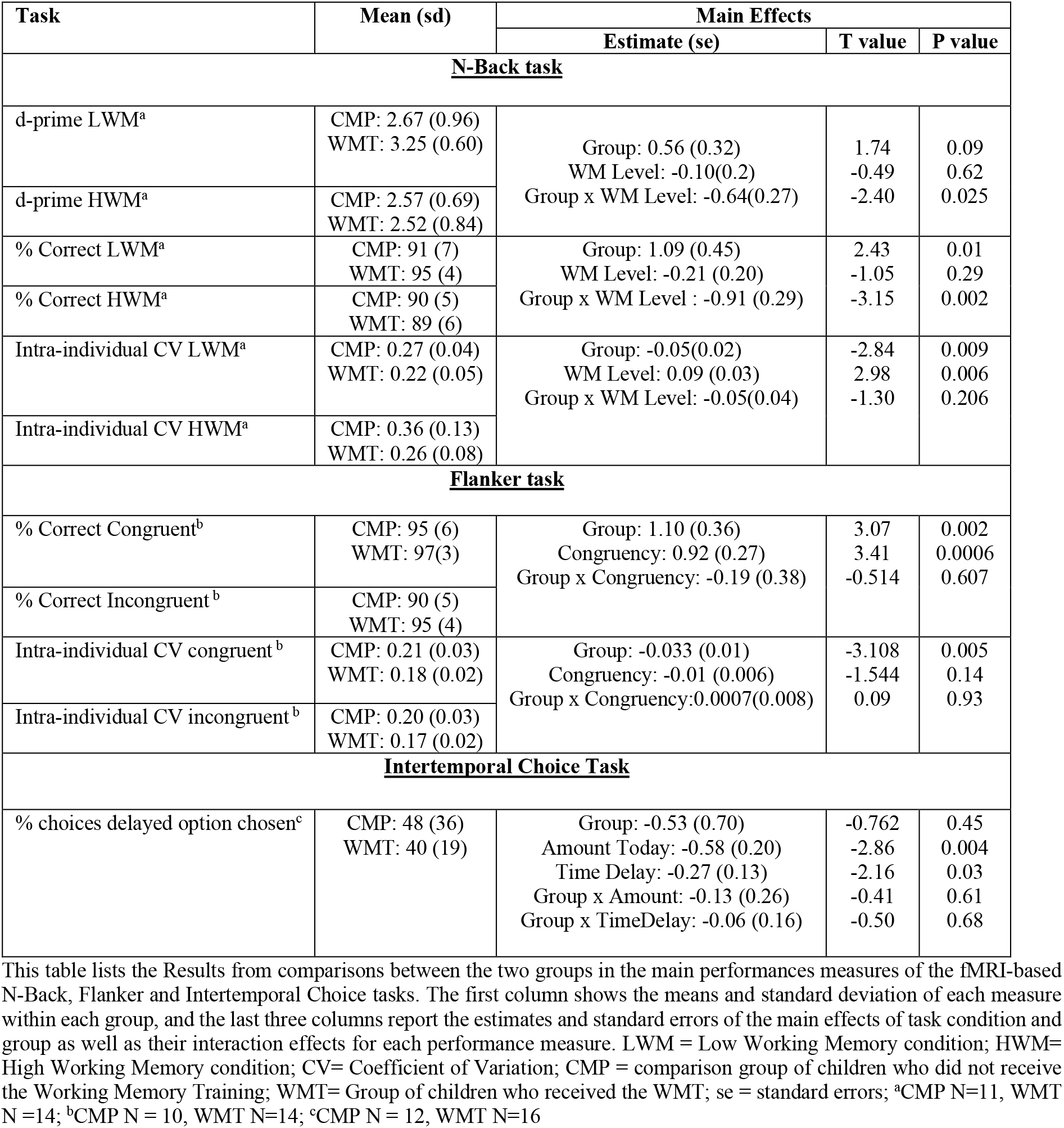
Results from group comparison in cognitive task performance measures.

**Table S3.** Between-groups differences in Ex-Gaussian reaction times.

| Ex-Gaussian parameter | Estimate of Group difference (SE) | T value | P value |
| --- | --- | --- | --- |
| Mu | -0.019 (0.022) | -0.851 | 0.4047 |
| Sigma | -0.020 (0.006) | -3.450 | 0.0025 |
| Tau | -0.021 (0.009) | -2.427 | 0.0248 |

This table shows the results from the analysis of ex-gaussian parameters fit to individual's response times. Group differences are computed as WMT – CMP groups. The T and P-values were computed from linear regressions explaining the variability in one parameter as a function of training group and the other two parameters of the ex-gaussian function (e.g. $\mu \sim 1 + \sigma + \tau + \text{Group}$ ).

**Table S4.** Group differences in ICV in DDM-simulated Flanker task response times.

|  | Estimate | Std.Error | T-Value | P Value |
| --- | --- | --- | --- | --- |
| Group | -0.108625 | 0.04441 | -2.446 | 0.0185 |
| Trial Type | 0.014553 | 0.047969 | 0.303 | 0.763 |
| Group x Trial Type | -0.005607 | 0.062806 | -0.089 | 0.9293 |

This table shows the results of a linear regression analysis testing whether or not the fitted DDM parameters can reproduce the patterns of response time variability seen in the two empirical groups. The dependent variable in this regression is the intra-individual coefficient of variation in response times (ICV) from simulated agents performing the Flanker task. Each simulated agent performed the task using the best-fitting parameters for the empirical data from one child. Analogous to the empirical data, we analyzed these simulated data as a function of Trial Type (Congruent, Incongruent) and Group (i.e. the training group the child whose parameters the agent used belonged to). The response times generated by the DDM simulations reproduce the group difference in ICV that we observe in the empirical data.

## Supplementary text

Here we give a brief description of the relevant measures used to assess the impact of the training are provided below. For an extensive description see Berger et al (2020).

Computerized stimuli were presented via headphones (when auditive) or on a touch-screen (when visual). Their responses had to be provided via touch-screen. All test scores were standardized to mean = 0 and SD = 1.

### 1. Working memory measures

Three different measures were used to identify the potential presence of transfer effects to non-trained working memory tests (near transfer effects).

‐ Verbal simple span. After listening to a series of one-digit numbers, the child must enter the sequence in the correct order on the touch-screen in front of him. Scores are calculated taking into account the number of correct sequences and the number of items in each sequence.
‐ Verbal complex span. This is a two-step task. In the first step, the child listens to a sequence of words describing objects, after each of them (s)he must indicate whether the object is an animal. After the sequence is finished, a grid with 3×3 images is shown, and the child has to reproduce the sequence of words heard in the correct order. As above, scores are calculated taking into account the number of correct sequences and the number of items in each sequence.
‐ Visuo-spatial complex span. This is also a two-step task. In the first step, the child is shown a series of 3 shapes in a single row and has to indicate the one that is different (left/center/right). After a number of screens, each with the three shapes shown in a row, the child has to indicate in an empty screen the position of the different stimulus on each screen, in the right sequence order. The difficulty increases by the increasing number of screens shown until the child gets to the empty response screen, thus increasing working memory demands with an increasingly longer sequence of positions.

### 2. Educational achievements

As above, three different measures were used to assess educational achievements, targeting reading, arithmetic and geometry skills. These assessments increased in difficulty with each follow-up assessment to incorporate the natural development of these skills throughout the school year.

‐ Reading: the child is presented with a series of sentences including a gap in the upper part of the screen. The child must fill in the gap choosing from the four alternative options presented in the lower part of the screen.
‐ Arithmetic: the final arithmetic score was the result of adding three different subscores. The first subscore was given by the “number sense” test, where participants are presented a varying number of circles/balls in a horizontal empty grid (2 rows x10 columns), and they have to compute the number of balls shown on each trial. While there are different mechanisms by which the child might get to the correct answer, the short presentation time (1.7s) prevents them from simply counting the balls and requires some basic arithmetic skills. The second subscore is obtained from the auditory arithmetic task, where the child listens to additions or subtractions of two numbers and has to provide the correct answer in the touch-screen. In the final written arithmetic subtest, the participant is presented with visual stimuli including at least three elements and they have to perform the calculation (addition/subtraction) sequentially to arrive to the correct solution, which has to be provided again with the touch screen.
‐ Geometry: the child is visually presented with a large geometric object and must determine how many smaller, simple-shaped object (i.e. square, rectangle, triangle) would fit into that larger object. Both are simultaneously presented on the screen and remain on screen until response is provided.

### 3. IQ measures

Two different sets of 17 items from the Colored Progressive Raven’s Matrices were used in an alternate fashion to assess IQ on each assessment wave.

## Footnotes

1 Cogmed and Cogmed Working Memory Training are trademarks, in the U.S. and/or other countries, of Cogmed Inc. (www.cogmed.com).

## Notes

### Competing Interest Statement

The authors have declared no competing interest.

## References

1 Schwaighofer, M., Fischer, F. & Buhner, M. Does Working Memory Training Transfer? A Meta-Analysis Including Training Conditions as Moderators. Educ Psychol 50, 138–166, (2015).

2 Melby-Lervag, M., Redick, T. S. & Hulme, C. Working Memory Training Does Not Improve Performance on Measures of Intelligence or Other Measures of “Far Transfer”: Evidence From a Meta-Analytic Review. Perspect Psychol Sci 11, 512–534, (2016).

3 Au, J. et al. Improving fluid intelligence with training on working memory: a meta-analysis. Psychon Bull Rev 22, 366–377, (2015).

4 Karbach, J. & Verhaeghen, P. Making working memory work: a meta-analysis of executive-control and working memory training in older adults. Psychological science 25, 2027–2037, (2014).

5 Cortese, S. et al. Cognitive training for attention-deficit/hyperactivity disorder: meta-analysis of clinical and neuropsychological outcomes from randomized controlled trials. Journal of the American Academy of Child and Adolescent Psychiatry 54, 164–174, (2015).

6 Wass, S. V., Scerif, G. & Johnson, M. H. Training attentional control and working memory - Is younger, better? Developmental Review 32, 360–387, (2012).

7 Bogg, T. & Lasecki, L. Reliable gains? Evidence for substantially underpowered designs in studies of working memory training transfer to fluid intelligence. Front Psychol 5, 1589, (2014).

8 Sala, G. & Gobet, F. Working memory training in typically developing children: A multilevel meta-analysis. Psychon Bull Rev 27, 423–434, (2020).

9 Smid, C. R., Karbach, J. & Steinbeis, N. Toward a Science of Effective Cognitive Training. Current Directions in Psychological Science 0, 0963721420951599, (2020).

10 Berger, E. M., Fehr, E., Hermes, H., Schunk, D. & Winkel, K. The Impact of Working Memory Training on Children’s Cognitive and Noncognitive Skills. NHH Dept. of Economics Discussion Paper, (2020).

11 Buschkuehl, M., Jaeggi, S. M. & Jonides, J. Neuronal effects following working memory training. Developmental cognitive neuroscience2 Suppl 1, S167–179, (2012).

12 Salmi, J., Nyberg, L. & Laine, M. Working memory training mostly engages general-purpose large-scale networks for learning. Neuroscience and biobehavioral reviews 93, 108–122, (2018).

13 Schneiders, J. A. et al. The impact of auditory working memory training on the fronto-parietal working memory network. Frontiers in human neuroscience 6, 173, (2012).

14 Flegal, K. E., Ragland, J. D. & Ranganath, C. Adaptive task difficulty influences neural plasticity and transfer of training. NeuroImage 188, 111–121, (2019).

15 Klingberg, T. Training and plasticity of working memory. Trends in cognitive sciences 14, 317–324, (2010).

16 McNab, F. et al. Changes in cortical dopamine D1 receptor binding associated with cognitive training. Science 323, 800–802, (2009).

17 D’Esposito, M. & Postle, B. R. The Cognitive Neuroscience of Working Memory. Annual Review of Psychology, Vol 66 66, 115–142, (2015).

18 Frank, M. J., Loughry, B. & O’Reilly, R. C. Interactions between frontal cortex and basal ganglia in working memory: A computational model. Cogn Affect Behav Ne 1, 137–160, (2001).

19 Mcnab, F. & Klingberg, T. Prefrontal cortex and basal ganglia control access to working memory. Nature neuroscience 11, 103–107, (2008).

20 Owen, A. M., McMillan, K. M., Laird, A. R. & Bullmore, E. N-back working memory paradigm: a meta-analysis of normative functional neuroimaging studies. Human brain mapping 25, 46–59, (2005).

21 Wager, T. D. & Smith, E. E. Neuroimaging studies of working memory: a meta-analysis. Cognitive, affective & behavioral neuroscience 3, 255–274, (2003).

22 Norman, D. A. & Shallice, T. Attention to Action - Willed and Automatic-Control of Behavior. B Psychonomic Soc 21, 354–354, (1983).

23 Bunge, S. A. & Wright, S. B. Neurodevelopmental changes in working memory and cognitive control. Current opinion in neurobiology 17, 243–250, (2007).

24 Darki, F. & Klingberg, T. The Role of Fronto-Parietal and Fronto-Striatal Networks in the Development of Working Memory: A Longitudinal Study. Cereb Cortex 25, 1587–1595, (2015).

25 Crone, E. A., Wendelken, C., Donohue, S., van Leijenhorst, L. & Bunge, S. A. Neurocognitive development of the ability to manipulate information in working memory. Proceedings of the National Academy of Sciences of the United States of America 103, 9315–9320, (2006).

26 Geier, C. F., Garver, K., Terwilliger, R. & Luna, B. Development of working memory maintenance. J Neurophysiol 101, 84–99, (2009).

27 Tamnes, C. K. et al. Longitudinal working memory development is related to structural maturation of frontal and parietal cortices. Journal of cognitive neuroscience 25, 1611–1623, (2013).

28 Montez, D. F., Calabro, F. J. & Luna, B. The expression of established cognitive brain states stabilizes with working memory development. Elife 6, (2017).

29 Matthaus, F. et al. Effects of age on the structure of functional connectivity networks during episodic and working memory demand. Brain Connect 2, 113–124, (2012).

30 Rypma, B., Prabhakaran, V., Desmond, J. E. & Gabrieli, J. D. Age differences in prefrontal cortical activity in working memory. Psychol Aging 16, 371–384, (2001).

31 Mattay, V. S. et al. Neurophysiological correlates of age-related changes in working memory capacity. Neurosci Lett 392, 32–37, (2006).

32 Dahlin, E., Neely, A. S., Larsson, A., Backman, L. & Nyberg, L. Transfer of learning after updating training mediated by the striatum. Science 320, 1510–1512, (2008).

33 Morrison, A. B. & Chein, J. M. Does working memory training work? The promise and challenges of enhancing cognition by training working memory. Psychon Bull Rev 18, 46–60, (2011).

34 MacDonald, S. W., Li, S. C. & Backman, L. Neural underpinnings of within-person variability in cognitive functioning. Psychol Aging 24, 792–808, (2009).

35 Saville, C. W. N. et al. On the stability of instability: Optimising the reliability of intra-subject variability of reaction times. Personality and Individual Differences 51, 148–153, (2011).

36 Kofler, M. J. et al. Reaction time variability in ADHD: A meta-analytic review of 319 studies. Clin Psychol Rev 33, 795–811, (2013).

37 Castellanos, F. X. et al. Varieties of attention-deficit/hyperactivity disorder-related intra-individual variability. Biological psychiatry 57, 1416–1423, (2005).

38 MacDonald, S. W., Nyberg, L. & Backman, L. Intra-individual variability in behavior: links to brain structure, neurotransmission and neuronal activity. Trends Neurosci 29, 474–480, (2006).

39 Williams, B. R., Hultsch, D. F., Strauss, E. H., Hunter, M. A. & Tannock, R. Inconsistency in reaction time across the life span. Neuropsychology 19, 88–96, (2005).

40 Papenberg, G., Hammerer, D., Muller, V., Lindenberger, U. & Li, S. C. Lower theta inter-trial phase coherence during performance monitoring is related to higher reaction time variability: a lifespan study. NeuroImage 83, 912–920, (2013).

41 Tamnes, C. K., Fjell, A. M., Westlye, L. T., Ostby, Y. & Walhovd, K. B. Becoming Consistent: Developmental Reductions in Intraindividual Variability in Reaction Time Are Related to White Matter Integrity. Journal of Neuroscience 32, 972–982, (2012).

42 van Belle, J. et al. Capturing the dynamics of response variability in the brain in ADHD. Neuroimage Clin 7, 132–141, (2015).

43 Johnson, B. P. et al. Left anterior cingulate activity predicts intra-individual reaction time variability in healthy adults. Neuropsychologia 72, 22–26, (2015).

44 Ilg, L., Klados, M., Alexander, N., Kirschbaum, C. & Li, S. C. Long-term impacts of prenatal synthetic glucocorticoids exposure on functional brain correlates of cognitive monitoring in adolescence. Sci Rep 8, 7715, (2018).

45 Bellgrove, M. A., Hester, R. & Garavan, H. The functional neuroanatomical correlates of response variability: evidence from a response inhibition task. Neuropsychologia 42, 1910–1916, (2004).

46 Isbell, E., Calkins, S. D., Swingler, M. M. & Leerkes, E. M. Attentional fluctuations in preschoolers: Direct and indirect relations with task accuracy, academic readiness, and school performance. J Exp Child Psychol 167, 388–403, (2018).

47 Montez, D. F., Calabro, F. J. & Luna, B. Working memory improves developmentally as neural processes stabilize. Plos One 14, e0213010, (2019).

48 Duckworth, A. L., Kirby, T., Gollwitzer, A. & Oettingen, G. From Fantasy to Action: Mental Contrasting with Implementation Intentions (MCII) Improves Academic Performance in Children. Soc Psychol Personal Sci 4, 745–753, (2013).

49 Rueda, M. R. et al. Development of attentional networks in childhood. Neuropsychologia 42, 1029–1040, (2004).

50 Steinbeis, N., Haushofer, J., Fehr, E. & Singer, T. Development of Behavioral Control and Associated vmPFC-DLPFC Connectivity Explains Children’s Increased Resistance to Temptation in Intertemporal Choice. Cereb Cortex, (2014).

51 RStudio Team. (2020).

52 Plummer, M. in Proceedings of the 3rd international workshop on distributed statistical computing. 1–10 (Vienna, Austria).

53 Wabersich, D. & Vandekerckhove, J. Extending JAGS: a tutorial on adding custom distributions to JAGS (with a diffusion model example). Behav Res Methods 46, 15–28, (2014).

54 Plummer, M. & Stukalov, A. Package “rjags’ [Software update for JAGS], 2018).

55 Woerner, W. et al. Normative data and evaluation of the German parent-rated Strengths and Difficulties Questionnaire (SDQ): Results of a representative field study. Z Kinder Jug-Psych 30, 105–112, (2002).

56 van Buuren, S. & Groothuis-Oudshoorn, K. mice: Multivariate imputation by chained equations in R. Journal of statistical software, 1–68, (2010).

57 Bürkner, P.-C. Advanced Bayesian Multilevel Modeling with the R Package brms. The R Journal 10, 395–411, (2018).

58 Stan Development Team. RStan: the R interface to Stan. R package version 2.18.1, (2018).

59 Achenbach, T. M. & Edelbrock, C. S. Manual for the child behavior checklist and revised child behavior profile. (1983).

60 Forstmann, B. U., Ratcliff, R. & Wagenmakers, E. J. Sequential Sampling Models in Cognitive Neuroscience: Advantages, Applications, and Extensions. Annual Review of Psychology, Vol 67 67, 641–666, (2016).

61 Ratcliff, R., Smith, P. L., Brown, S. D. & McKoon, G. Diffusion Decision Model: Current Issues and History. Trends in cognitive sciences 20, 260–281, (2016).

62 Cavanagh, J. F., Wiecki, T. V., Kochar, A. & Frank, M. J. Eye tracking and pupillometry are indicators of dissociable latent decision processes. J Exp Psychol Gen 143, 1476–1488, (2014).

63 Krajbich, I., Hare, T., Bartling, B., Morishima, Y. & Fehr, E. A Common Mechanism Underlying Food Choice and Social Decisions. PLoS computational biology 11, e1004371, (2015).

64 Krajbich, I. & Rangel, A. Multialternative drift-diffusion model predicts the relationship between visual fixations and choice in value-based decisions. Proceedings of the National Academy of Sciences of the United States of America 108, 13852–13857, (2011).

65 Casey, B. J. et al. The Adolescent Brain Cognitive Development (ABCD) study: Imaging acquisition across 21 sites. Developmental cognitive neuroscience 32, 43–54, (2018).

66 Unsworth, N. & Robison, M. K. A locus coeruleus-norepinephrine account of individual differences in working memory capacity and attention control. Psychon Bull Rev 24, 1282–1311, (2017).

67 Eriksson, J., Vogel, E. K., Lansner, A., Bergstrom, F. & Nyberg, L. Neurocognitive Architecture of Working Memory. Neuron 88, 33–46, (2015).

68 Astle, D. E. & Scerif, G. Interactions between attention and visual short-term memory (VSTM): what can be learnt from individual and developmental differences? Neuropsychologia 49, 1435–1445, (2011).

69 Baddeley, A. in Attention: Selection, awareness, and control (eds A. Baddeley & L. Weiskrantz) 152–170 (Oxford University Press, 1993).

70 Engle, R. W. Working Memory and Executive Attention: A Revisit. Perspect Psychol Sci 13, 190–193, (2018).

71 Gazzaley, A. & Nobre, A. C. Top-down modulation: bridging selective attention and working memory. Trends in cognitive sciences 16, 129–135, (2012).

72 Zanto, T. P., Rubens, M. T., Thangavel, A. & Gazzaley, A. Causal role of the prefrontal cortex in top-down modulation of visual processing and working memory. Nature neuroscience 14, 656–661, (2011).

73 Spencer-Smith, M. & Klingberg, T. Benefits of a Working Memory Training Program for Inattention in Daily Life: A Systematic Review and Meta-Analysis. PloS one 10, (2015).

74 Greenwood, P. M. & Parasuraman, R. The mechanisms of far transfer from cognitive training: Review and hypothesis. Neuropsychology 30, 742–755, (2016).

75 Fukuda, K. & Vogel, E. K. Individual Differences in Recovery Time From Attentional Capture. Psychological science 22, 361–368, (2011).

76 Stuss, D. T., Murphy, K. J., Binns, M. A. & Alexander, M. P. Staying on the job: the frontal lobes control individual performance variability. Brain : a journal of neurology 126, 2363–2380, (2003).

77 Unsworth, N. Consistency of attentional control as an important cognitive trait: A latent variable analysis. Intelligence 49, 110–128, (2015).

78 Kelly, A. M., Uddin, L. Q., Biswal, B. B., Castellanos, F. X. & Milham, M. P. Competition between functional brain networks mediates behavioral variability. NeuroImage 39, 527–537, (2008).

79 Larson, G. E. & Saccuzzo, D. P. Cognitive Correlates of General Intelligence -toward a Process Theory of G. Intelligence 13, 5–31, (1989).

80 Walhovd, K. B. & Fjell, A. M. White matter volume predicts reaction time instability. Neuropsychologia 45, 2277–2284, (2007).

81 Li, S.-C. et al. Transformations in the Couplings Among Intellectual Abilities and Constituent Cognitive Processes Across the Life Span. Psychological science 15, 155–163, (2004).

82 Karalunas, S. L., Geurts, H. M., Konrad, K., Bender, S. & Nigg, J. T. Annual research review: Reaction time variability in ADHD and autism spectrum disorders: measurement and mechanisms of a proposed trans-diagnostic phenotype. Journal of child psychology and psychiatry, and allied disciplines 55, 685–710, (2014).

83 Geurts, H. M. et al. Intra-individual variability in ADHD, autism spectrum disorders and Tourette’s syndrome. Neuropsychologia 46, 3030–3041, (2008).

84 Haynes, B., Bauermeister, S. & Bunce, D. A systematic review of longitudinal associations between reaction time intraindividual variability and age-related cognitive decline or impairment, dementia, and mortality. Journal of the International Neuropsychological Society 23, 431–445, (2017).

85 Kuntsi, J. et al. Separation of Cognitive Impairments in Attention-Deficit/Hyperactivity Disorder Into 2 Familial Factors. Archives of general psychiatry 67, 1159–1167, (2010).

86 Adleman, N. E. et al. Increased intrasubject variability in response time in unaffected preschoolers at familial risk for bipolar disorder. Psychiatry Res 219, 687–689, (2014).

87 van Ravenzwaaij, D., Brown, S. & Wagenmakers, E. J. An integrated perspective on the relation between response speed and intelligence. Cognition 119, 381–393, (2011).

88 McKenzie, D. Beyond baseline and follow-up: The case for more T in experiments. Journal of Development Economics 99, 210–221, (2012).

89 Karbach, J., Strobach, T. & Schubert, T. Adaptive working-memory training benefits reading, but not mathematics in middle childhood. Child Neuropsychology 21, 285–301, (2015).

90 Titz, C. & Karbach, J. Working memory and executive functions: effects of training on academic achievement. Psychological research 78, 852–868, (2014).

